# DNA amplification in mixed aqueous-organic media: chemical analysis of leading polymerase chain reaction compositions

**DOI:** 10.1101/2021.10.02.462877

**Authors:** Jalel Neffati, Ioanna Petrounia, Rudy D. Moreira, Raj Chakrabarti

## Abstract

PCR amplification of GC-rich regions often leads to low yield and specificity. Addition of PCR-enhancing compounds is employed in order to overcome these obstacles. PCR-enhancing additives are low molecular polar organic compounds that are included as undisclosed co-solvents in commercial PCR buffers. In the interest of transparency and to permit further optimization by researchers of PCR compositions for challenging amplification problems, we studied eight PCR buffers by GC/MS to identify and quantify their co-solvents. Buffer specificity, both rich in water and salified substances, required a suitable sample preparation before injection into the GC/MS system. The aqueous phase of each buffer was replaced by an organic solvent to remove, by precipitation and filtration, salified substances which are detrimental to the GC/MS analysis. This approach has demonstrated the advantage of eliminating both water and salified substances without any loss of co-solvents. The sensitivity of the developed method was demonstrated as the main co-solvents were easily detected, identified and quantified. The methodology for identifying the co-solvents is mainly based on comparison of both library matching of acquired MS spectra with NIST library and experimental mass spectra obtained from authentic chemical standards. For the quantification of each co-solvent, deuterated Internal standards of similar structure to the cosolvents were used to correct the variable recovery caused by sample preparation, matrix effects, and ion source variability. The recovery ratio of the developed method was verified and found to be in the range 90-120 %. We then characterized the effects of specific organic co-solvents identified during PCR amplification -- using DNA melting, polymerase thermostability, polymerase activity and real-time PCR methods -- in order to elucidate their mechanism of action and to permit further optimization of their effects on amplification efficiency and specificity.

## Introduction

PCR is widely used in diagnostic and molecular analysis of DNA and RNA. Amplification of templates with a high GC content using PCR is usually difficult compared to low GC targets. PCR amplification of GC-rich nucleotide sequences is often associated with insufficient yield of the target DNA sequence and amplification of non-specific products [1–5]. GC-rich templates are difficult to denature because of their high melting temperature. Primer extension is impeded as well due to their greater tendency to form secondary structures. 28% of the human genome sequences are categorized into GC-rich genes (>60% GC content). Moreover, many important regulatory domains including promoters, enhancers and other control elements consist of GC-rich regions [6]. A common strategy has been to include small quantities of PCR-enhancing compounds to improve GC-rich gene amplification. These co-solvents belong to the groups of amides, sulfones, sulfoxides and diols. PCR-enhancing compounds are not disclosed in commercially available PCR buffers.

In the interest of transparency, we developed a simple and robust analytical methodology to identify and quantify the co-solvents in eight commercial PCR buffers. We selected GC/MS over LC/MS for our studies as it appears to be the best tool for rapid analysis of these compounds [7–10]. Co-solvents are low molecular weight, thermally stable and sufficiently volatile. Another advantage of GC/MS is that this technique (by means of electron ionization mass spectrometry) allows easier and faster identification based on a search into reference libraries [11].

However, the GC/MS technique has a weakness with respect to the analysis of products containing non-volatile salified substances and/or high concentration of water [12–13]. Aqueous buffers are in this case as they contain many salified organic or inorganic (such as Tris.HCl, KCl, MgCl_2_) level of water. It was therefore decided to develop an analytical method with a pretreatment of the buffers before injection in the GC/MS system. The aqueous phase of each buffer was replaced with an organic solvent with the objective of precipitating the salts and filtering them. Specifically, the water was removed in a simple manner by lyophilization, then the lyophilizate was dissolved in a judiciously chosen organic diluent.

Even though co-solvents enhance amplification through their favorable effects on DNA duplex stability, they often compromise the thermostability and activity of the polymerase. We therefore evaluated the effect of select co-solvents on Taq polymerase and DNA melting.

## 1 Materials and Methods

### 1.1 PCR Buffers

Eight buffers were analyzed with the developed method. Table 1 gives the description of each kit with its buffer as well as the manufacturer and the commercial supplier. Three Toyobo buffers were purchased from Diagnocine (US). Two KAPA Biosystems buffers were purchased from Roche Diagnostic (France) and Aldrich (France). NEB and Sigma buffers were purchased from NEB (UK) and Sigma (France), respectively.

**Table 1:** Origin of eight commercial PCR Buffers

| PCR Kit | Manufacturer | Cat # (Supplier) | PCR Buffer |
| --- | --- | --- | --- |
| GC Rich PCR system | Sigma | 4743784001<br>(Aldrich) | PCR Reaction Buffer |
| KAPA HiFi PCR Kit | Kapa Biosystems | KK2103<br>(Roche diagnostics) | GC Buffer |
| KAPA2G Robust PCR Kit | Kapa Biosystems | KK5024<br>(Aldrich) | GC Buffer |
| Q5 High fidelity DNA polymerase | NEB | M0491S<br>(NEB) | Q5 High GC Enhancer |
| Thunderbird | Toyobo | QRZ-101<br>(Diagnocine) | 2X reaction buffer |
| Thunderbird Probe qPCR Mix | Toyobo | QPS-101<br>(Diagnocine) | Thunderbird Probe qPCR Mix |
| Thunderbird Probe qPCR Mix | Toyobo | QPS-201<br>(Diagnocine) | Thunderbird Probe qPCR Mix |
| KOD SYBR qPCR Mix | Toyobo | QKD-201<br>(Diagnocine) | KOD SYBR<sup>®</sup> qPCR Mix |

**Table 2:**
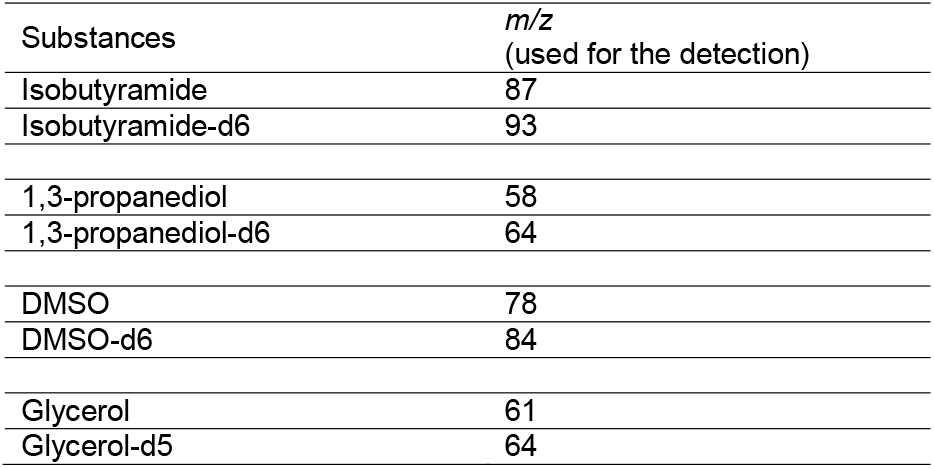
m/z Ion selected for the quantification of each co-solvent

| Substances | *m/z*<br>(used for the detection) |
| --- | --- |
| Isobutyramide | 87 |
| Isobutyramide-d6 | 93 |
| 1,3-propanediol | 58 |
| 1,3-propanediol-d6 | 64 |
| DMSO | 78 |
| DMSO-d6 | 84 |
| Glycerol | 61 |
| Glycerol-d5 | 64 |

### 1.2 Standards and deuterated standards

Co-solvent standards were used both for identification and quantification of the main compounds of interest. Standards of 1,3-propanediol (98 %, Cat # P50404-100G) and isobutyramide (99 %, Cat #144436-25G) were purchased from Sigma Aldrich (France). Dimethylsulfoxide (DMSO) and glycerol were purchased from Merck (France) (Emsure ACS, Cat #144436-25G and ≥99%, Cat # 8.18709.1000, respectively)

Deuterated standards used as Internal Standard (IS) were specifically purchased for the quantification of each co-solvent: 1,3-propanediol-d6 (98%, Cat # P760322) and isobutyramide-d6(98 %, Cat # M325939) were purchased from Toronto Research Chemical (Canada). DMSO-d6 (≥ 99 % Cat # 156914-1G) and glycerol-d5 (98 %, Cat # 454524-1G) were purchased from Sigma Aldrich (France).

### 1.3 Lyophilization conditions

The buffers were lyophilized as the direct injection of aqueous buffers tended to rapidly damage the stationary phase of the GC column. The removal of the water was achieved first after introducing the commercial buffer into a 50 ml flask immersed in a bath containing dry ice and acetone. Once the buffer is frozen, the flask is connected to a rotary vane pump which permits to reach around 0.5-1 mbar. After 3 hours, the lyophilized buffer is recovered by dilution into 1 ml of an appropriate diluent. Finally, this solution is filtered through a 0.45 μm PTFE membrane and injected into the GC/MS system.

The choice of lyophilized buffer diluent is an important part of the co-solvent analysis, as the buffers contain non-volatile substances (such as MgCl_2_, KCl, Tris,HCl) that cannot be introduced into the GC injector. We therefore searched for a diluent that solubilizes only cosolvents of interest and which can remove the salified substances by filtration through a 0.45 μm PTFE membrane. Three diluents were tested: DMSO, MeOH and THF. DMSO and THF were not retained; it was observed that in DMSO, salified substances were completely soluble. THF was not selected, even though the salts were not solubilized because after filtration and injection of the lyophilized buffer in the system we did not detect any of the substances of interest. Finally, methanol was chosen because it allowed us to both remove salts substances by simple filtration, and to properly analyze co-solvents of interest.

### 1.4 GC/MS conditions

The analysis was performed with a Thermo Scientific GC/MS system with triple quadrupole mass analyzer. Equipment consisted of a TriPlus AS autosampler (for injection of liquid or gas samples), and a Trace GC Ultra connected to a TSQ Quantum GC mass spectrometer. The control of the GC/MS system and data collection were done by XCalibur 2.1 Software. Identification of cosolvents from their mass spectra was done using the by NIST MS search 2.2 software.

The GC separation was conducted on a Phenomenex column ZB-WAX 20 cm x 0.18 mm and a film thickness of 0.18 μm. Helium was used as carrier gas at a constant flow of 1 mL/min. The GC oven temperature of 40°C was increased at 20°C/min to 250°C. The final temperature was held for 1 min. The inlet temperature was kept at 250°C in split mode at 10mL/min. A straight glass injection liner with glass wool was obtained from Restek (Topaz precision liner, Ref: 23327, France).

The sample solutions or the standard solutions were injected with a 1-μL injection volume. After chromatographic separation, the compounds were transferred toward the mass source with a transfer line maintained at 200°C.

The MS was operated with two Ionization modes: 70eV Electron Impact Ionization (EI) and Chemical Ionization (CI) with methane and ammonia reagents; each of them was used with an ionization current at 28 μA and source temperature at 250°C. For identification purposes, full scan of the components was conducted in the range of 50-500 m/z (60-500 m/z for CI). For quantification, the mass spectrometer for Single Ion Monitoring mode (SIM mode) was set as follows:

### 1.5 Preparation of sample and standard solutions

For identification purposes, the overall contents of a PCR buffer bottle were lyophilized, then the product was diluted into 1ml of methanol as described in section 1.3. In both identification modes used (EI or CI mode), 1 μL of the methanol solution was directly injected in the GC/MS system.

Specifically for the quantification of each co-solvent, two solutions were prepared according to the amount described in Table 3: the first sample solution contained both the buffer and the d-internal standard, the second standard solution contained both the co-solvent standard and the d-internal standard. The sample solution was lyophilized, recovered into 1 mL of methanol, filtered and then diluted in accordance with initial concentrations of the substance to be quantified to obtain a response around 3×10^7^ counts (in the linear range of the detector). The standard solutions were not lyophilized but only diluted into 50 mL volumetric flasks with methanol (except for DMSO which was diluted into 100mL volumetric flasks).

**Table 3:** Preparation of buffer and standard solution

| Substance to quantify | Sample solutions |  | Standard solution |  |
| --- | --- | --- | --- | --- |
|  | Weight of Buffer (mg) | Weight of D-IS (mg) | Co-solvent (mg) | Weight of D-IS (mg) |
| Isobutyramide | 500 | 15 | 30 | 15 |
| 1,3-Propanediol | 500 | 30 | 30 | 15 |
| DMSO | 500 | 40 | 15 | 15 |
| Glycerol | 500 | 30 | 30 | 30 |

### 1.6 c-Jun amplification

c-Jun amplification was carried out using primers ATGACTGCAAAGATGGAAACG and TCAAAATGTTTGCAACTGCTGCG. 1 ng c-Jun and 1 unit Taq polymerase were used in the experiment. The PCR protocol was as follows : 94°C for 3 min, 94°C for 45 sec, 55°C for 30 sec, 72°C for 90 sec, 72°C for 10 min; steps 2-4 for 25 cycles.

### 1.7 Polymerase thermostability RT-PCR assay

A real-time PCR SYBR Green I-based assay [14] was used as a polymerase thermostability screen by subjecting enzymes to high temperature prior to PCR amplification. ~70 million anhydrotetracycline-induced Taq cells resuspended in 10 μl 1x Taq buffer (10 mM Tris-HCl, pH 8.0, 50 mM KCl, 1.5 mM MgCl_2_, 0.1% Triton X-100) were added to 40 μl of of the qPCR mix. The qPCR mix contained the appropriate amount of co-solvent, 0.31 mM dNTP, 1.25 mg/ml BSA, 3.75 mM MgCl_2_, 0.62 μM of primers Q1 (GGTCACCCGTTCAACCTGAACAG) and Q2 (GTCAACCGCCTTCACGCGGAAC), 0.62x SYBR GreenI in 1x Taq buffer. (Q1 and Q2 anneal to the Taq gene and produce a 513 bp product.) The 96-well plates were sealed and read on a BioRad CFX Real-Time PCR instrument. The PCR protocol was: 95°C for 6 min, 94°C for 30 sec, 57.8°C for 30 sec, 72°C for 30 sec; steps 2-4 for 16 cycles. Melting curves were collected in the 55°C-95°C temperature range. Data were analyzed using the CFX Manager. For lysates (crude screening), prior to the assay, lysates were prepared by heating the resuspended cells at 95°C for 5 min. Peak area data were corrected for cell number to provide a screening score for thermostability.

The RT-PCR assay was also used to screen for thermostability of purified Taq protein (15 nM Taq).

### 1.8 Polymerase activity assays

#### RT-PCR-based screening assay

Two-step PCR in the presence or absence of co-solvents was performed as a polymerase enzyme activity screen by challenging enzymes to amplify the DNA template at the annealing temperature without an extension step at 72 °C. The assay was performed as above but with the aforementioned omission of an extension step and also with the omission of the 95°C for 6 min heating at the outset of PCR cycling. The PCR protocol was as follows: 95°C for 1 min, 94°C for 30 sec, 57.8°C for 5 sec; steps 2-3 for 20 cycles.

#### Specific activity assay

The activity of purified Taq was determined using the EvaEz fluorometric polymerase activity assay kit (Biotium). 0.4 nM Taq were used in the assay.

### 1.9 Polymerase thermal denaturation (half-life) assay

The PicoGreen-based fluorescence assay was used to determine thermostability [15,16]. The primer/template conjugate (PTC) solution was prepared as follows: the M13 universal primer (GTTTTCCCAGTCACGACG) was mixed with M13mp18 at a 10x ratio in 50 mM KCl, 10 mM Tris-HCl, 1.5 mM MgCl_2_, pH 8.0. The PTC solution was heated at 70°C for 5 min and slowly cooled at RT. For the PicoGreen assay, 5% (v/v) 1,4-butanediol was incubated with 200 nM purified His-tagged Taq polymerase. The sample was heated at select temperatures and time points and cooled immediately on ice. The solution was made 20 ng/μl in TPC and 200 μM in dNTPs in a total volume of 50 μl and heated to 72°C for 15 min. 98 μl TE buffer, 2 μl PCR reaction and 100 μl of 1:200 (v/v) PicoGreen were combined in a black 96-well plate. Following a 10-min incubation in the dark, the fluorescence was measured using excitation at 480 nm and emission at 525 nm on a TECAN infinite M200 Pro (gain: 50).

## 2 Results

### 2.1 Co-solvent identification

Applying our methodology, we identified co-solvents in eight commercial PCR buffers. To achieve this, mass spectra were compared to the NIST library data as a as a first step in the identification of possible candidates and ranked according to their similarity index. The definitive identification of co-solvents was based on a comparison with experimental mass spectra obtained from authentic chemical standards.

Table 4 provides the data for each buffer with the retention time (RT) of each detected compound and the main ions m/z by EI compared to the most likely spectrum given by the NIST library. Definitive identification was obtained both by determining the molecular mass by CI and comparing the data obtained (i.e, the RT, and m /z) with those obtained from authentic co-solvent standards.

**Table 4:**
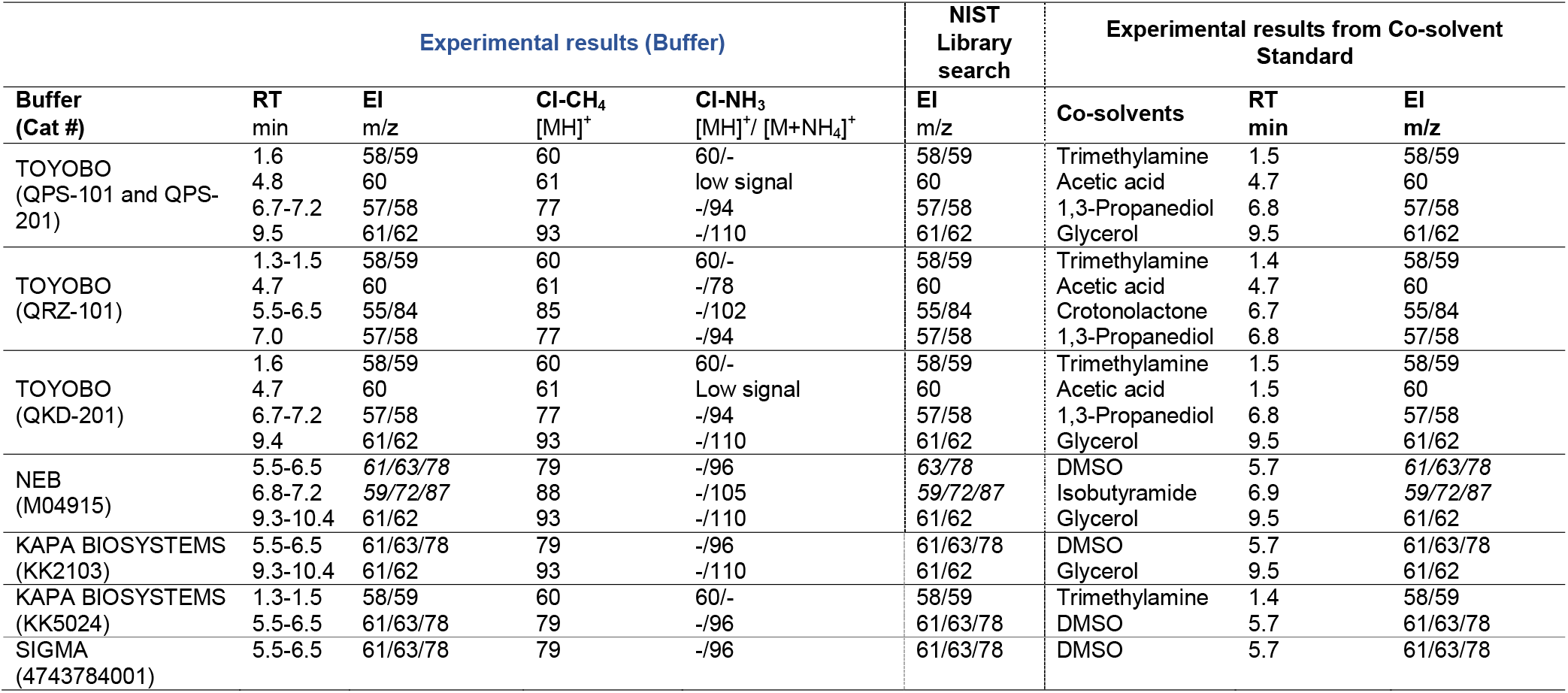
Identification by GC/MS-EI and CI of the co-solvents detected in seven PCR Buffers

Figures 1 to 7 illustrate the GC/MS-EI profile of all buffers. Figures 8 to 14, illustrate the EI-Mass spectra for each co-solvent detected in the NEB and Toyobo buffers compared to the mass spectra obtained from both the NIST library and co-solvent standard.

**Figure 1:**
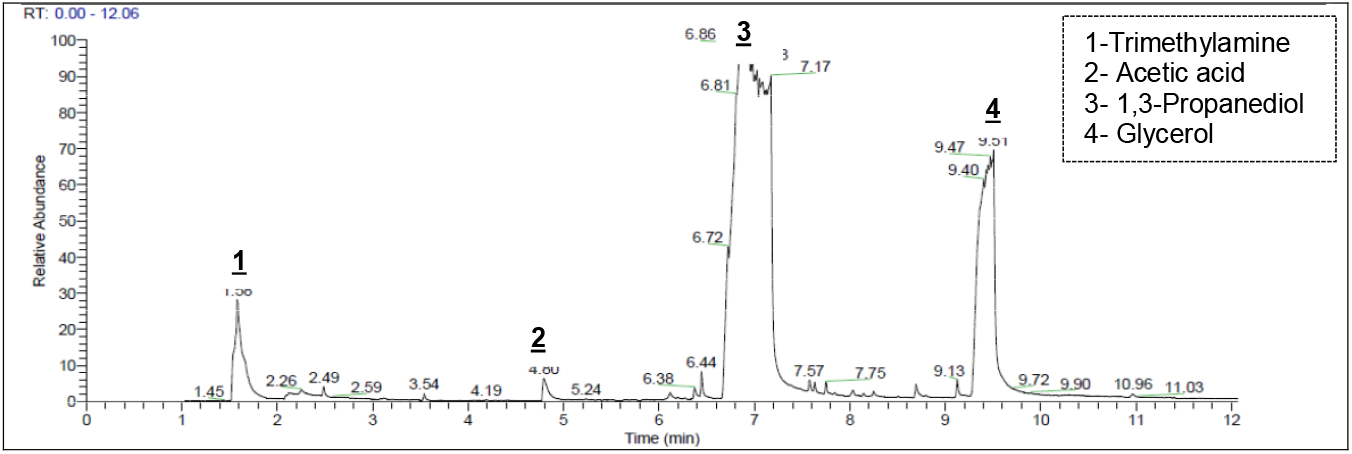
GC/MS chromatogram of Toyobo Cat # QPS-101 (similar to QPS-201)

**Figure 2:**
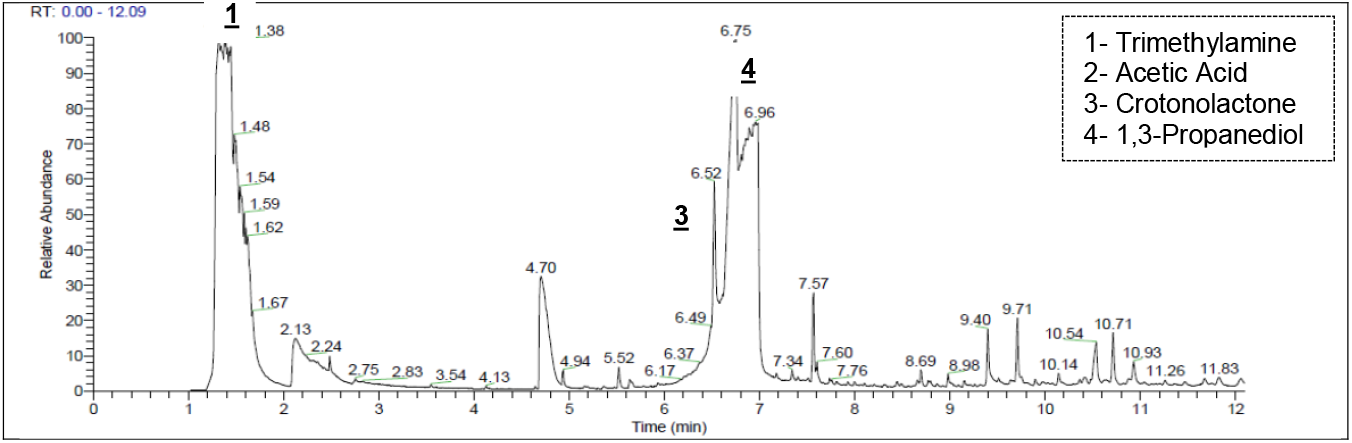
GC/MS chromatogram of Toyobo Cat # QRZ-101

**Figure 3:**
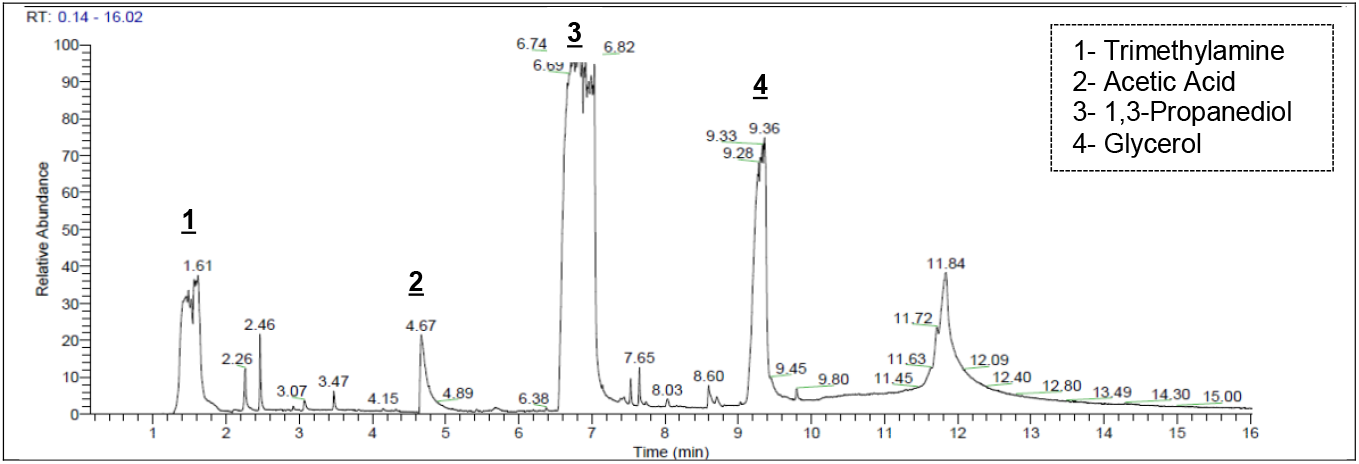
GC/MS chromatogram of Toyobo Cat # QKD 201

**Figure 4:**
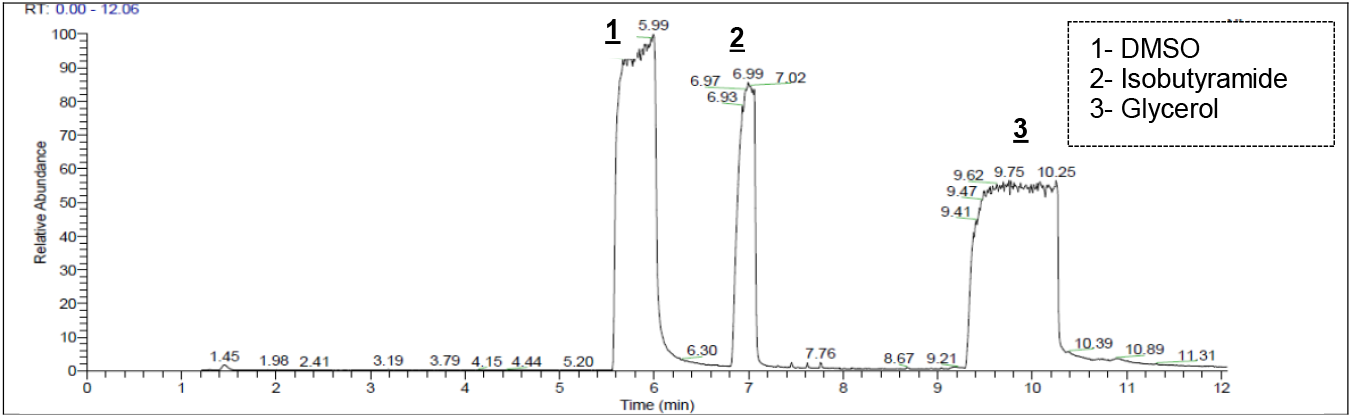
GC/MS chromatogram of NEB buffer Cat # M0491s

**Figure 5.**
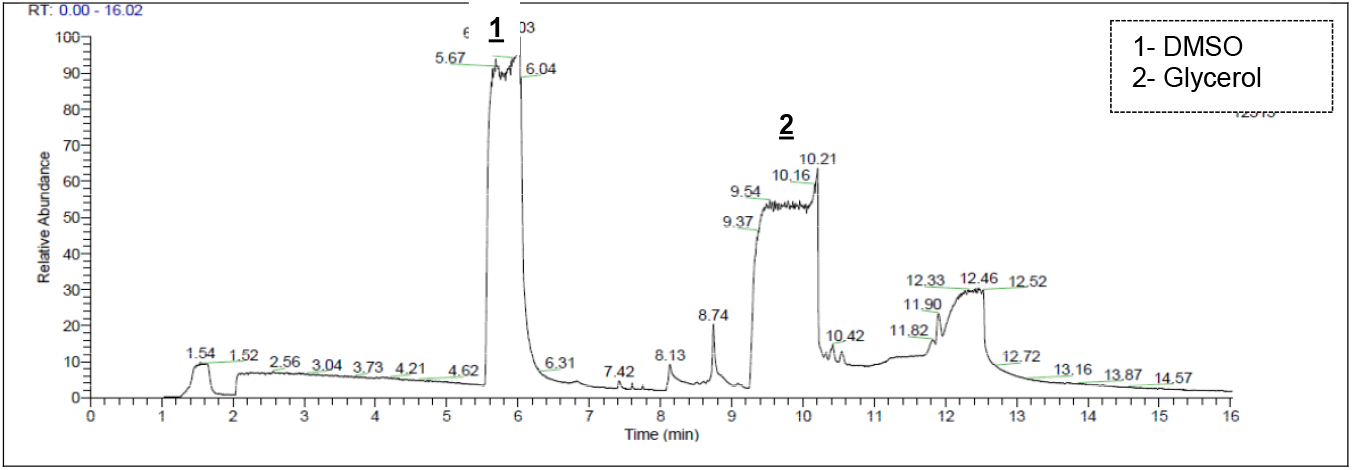
GC/MS chromatogram of Kapa Biosystems Buffer Cat # KK2103

**Figure 6:**
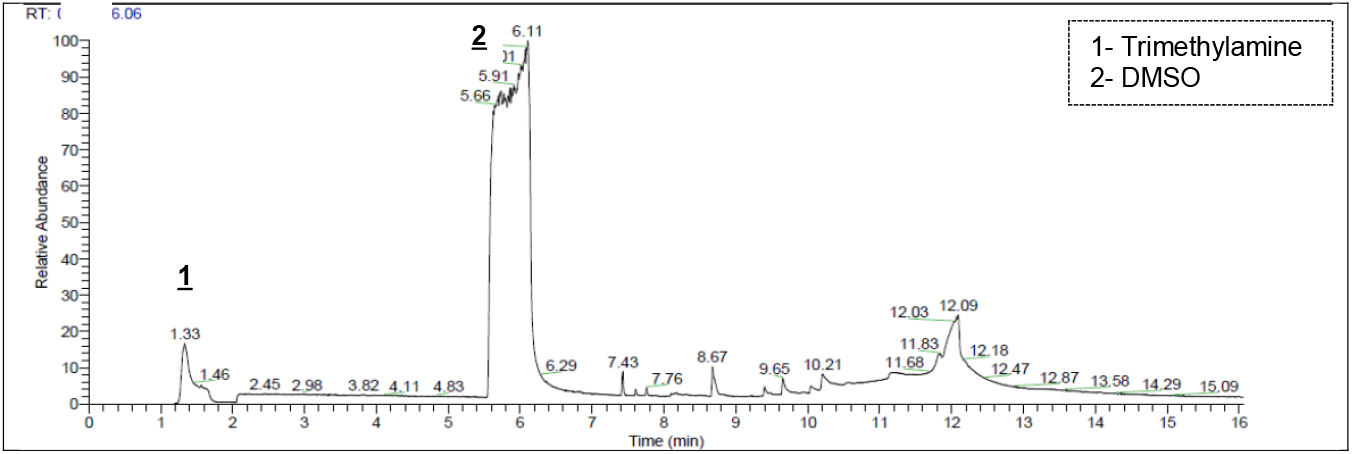
GC/MS chromatogram of Kapa Biosystems Buffer Cat # KK5024

**Figure 7:**
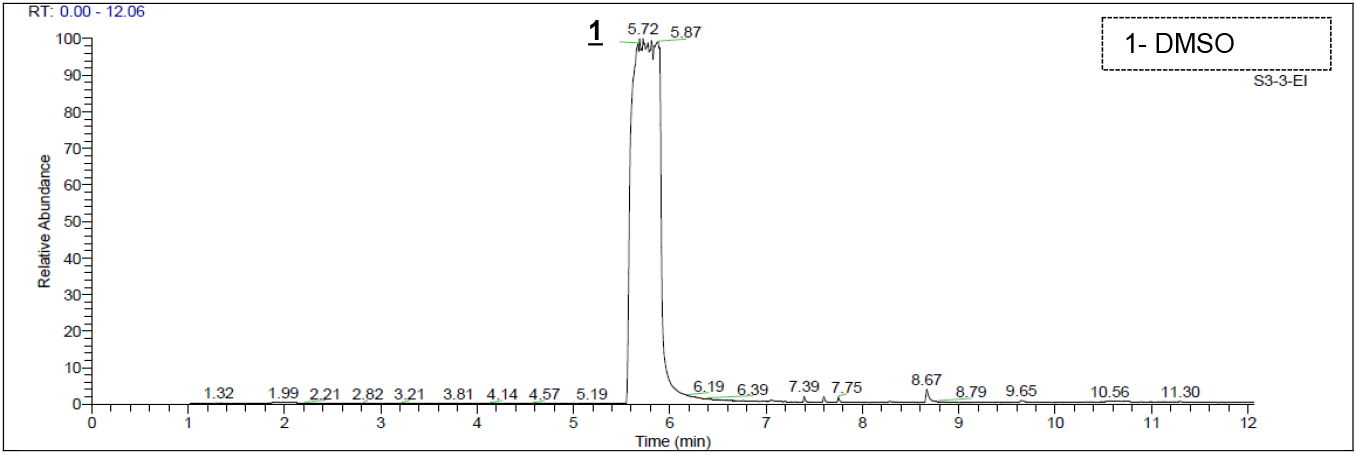
GC/MS chromatogram of Sigma buffer Cat # 4743784001

**Figure 8:**
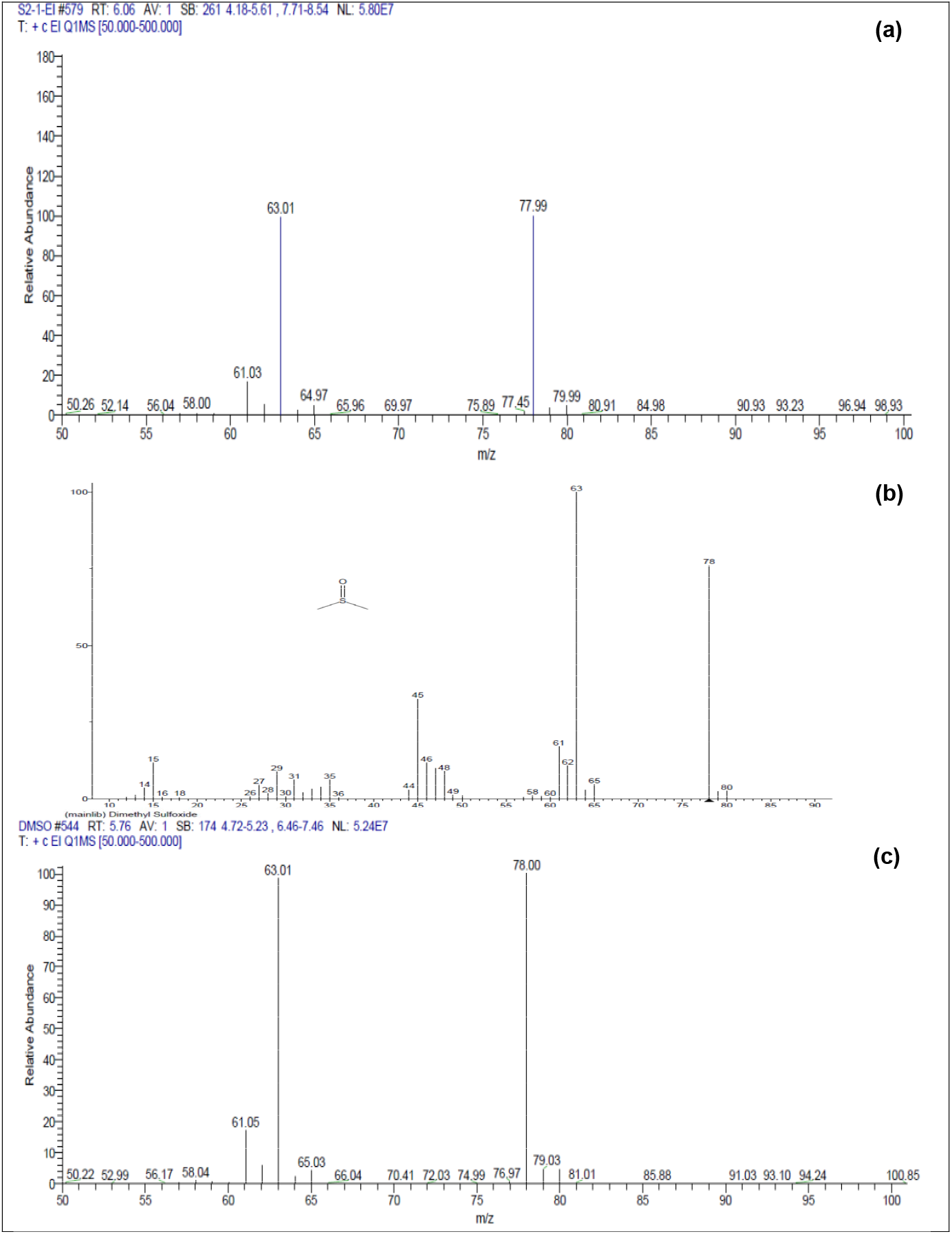
**(a)** EI-Mass spectra of DMSO at RT 6.0 min in the NEB buffer Cat # M0491s **(b)** EI-Mass spectra of DMSO given by the NIST Library **(c)** Experimental EI-Mass spectra of DMSO standard.

**Figure 9:**
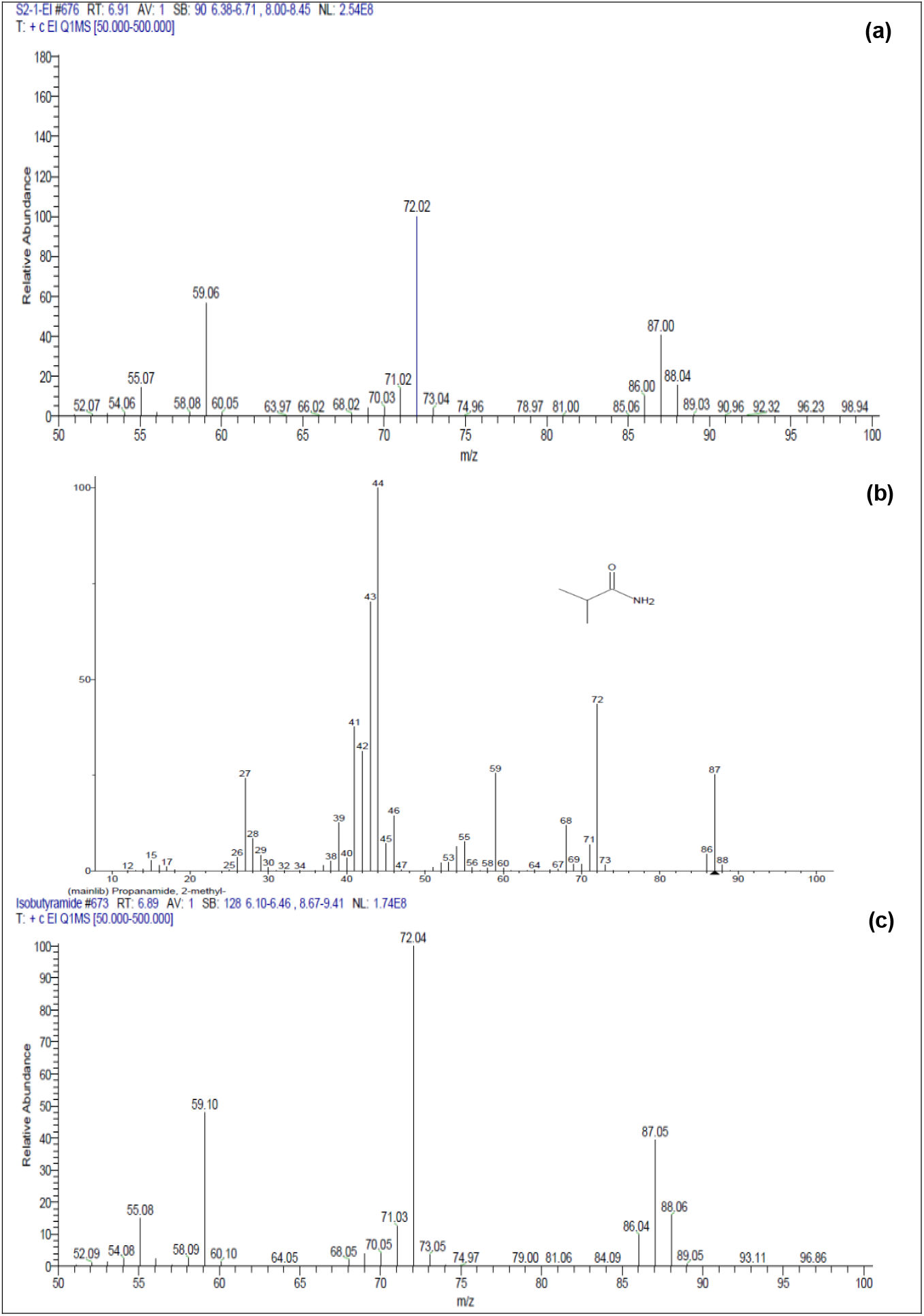
**(a)** EI-Mass spectra of isobutyramide at RT 7.0 min in the NEB buffer Cat # M0491s **(b)** EI-Mass spectra of isobutyramide given by the NIST Library **(c)** Experimental EI-Mass spectra of isobutyramide standard.

**Figure 10:**
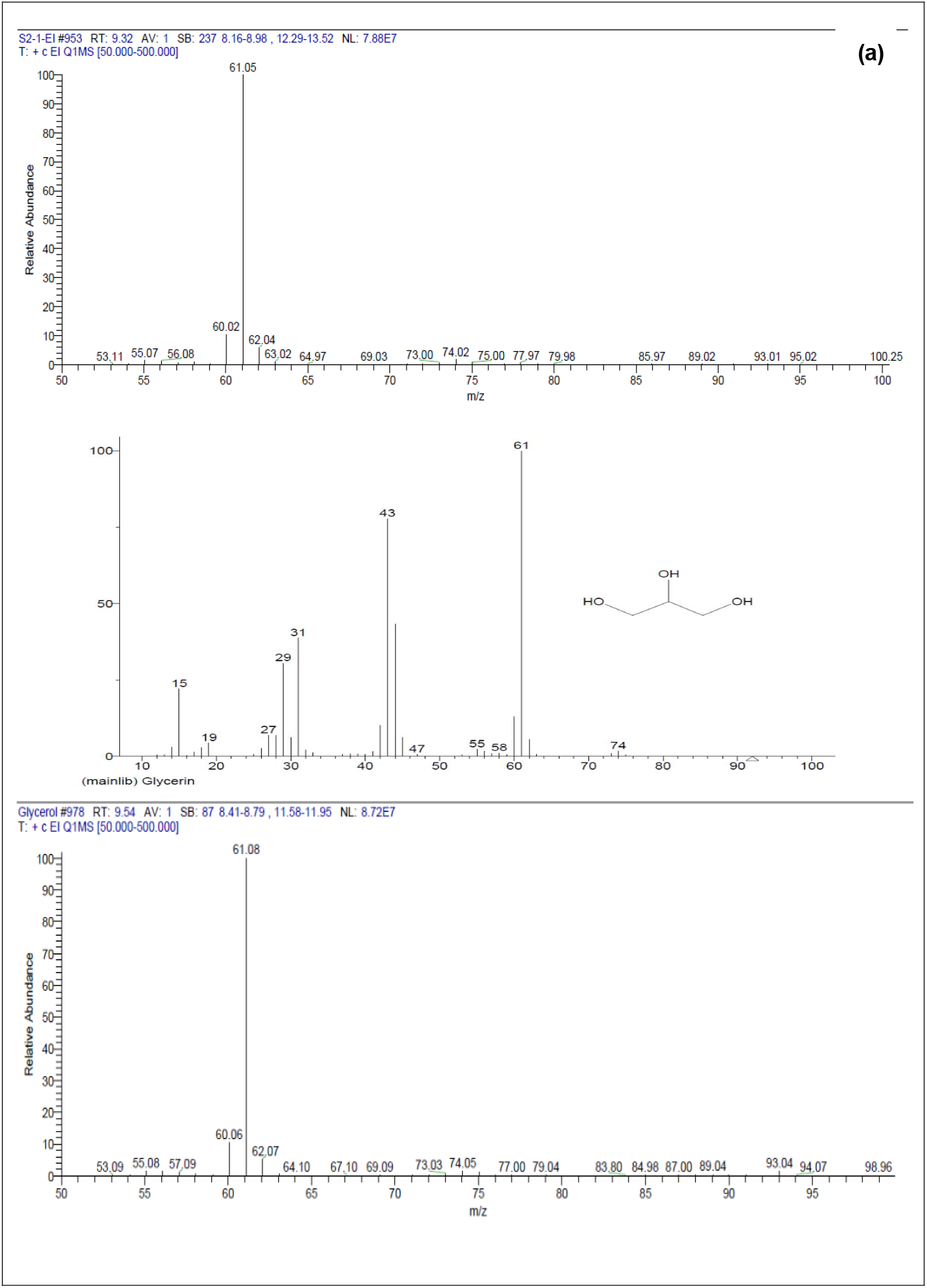
**(a)** EI-Mass spectra of glycerol at RT 9.8 min in the NEB buffer Cat # M0491s **(b)** EI-Mass spectra of glycerol given by the NIST Library **(c)** Experimental EI-Mass spectra of glycerol standard.

**Figure 11:**
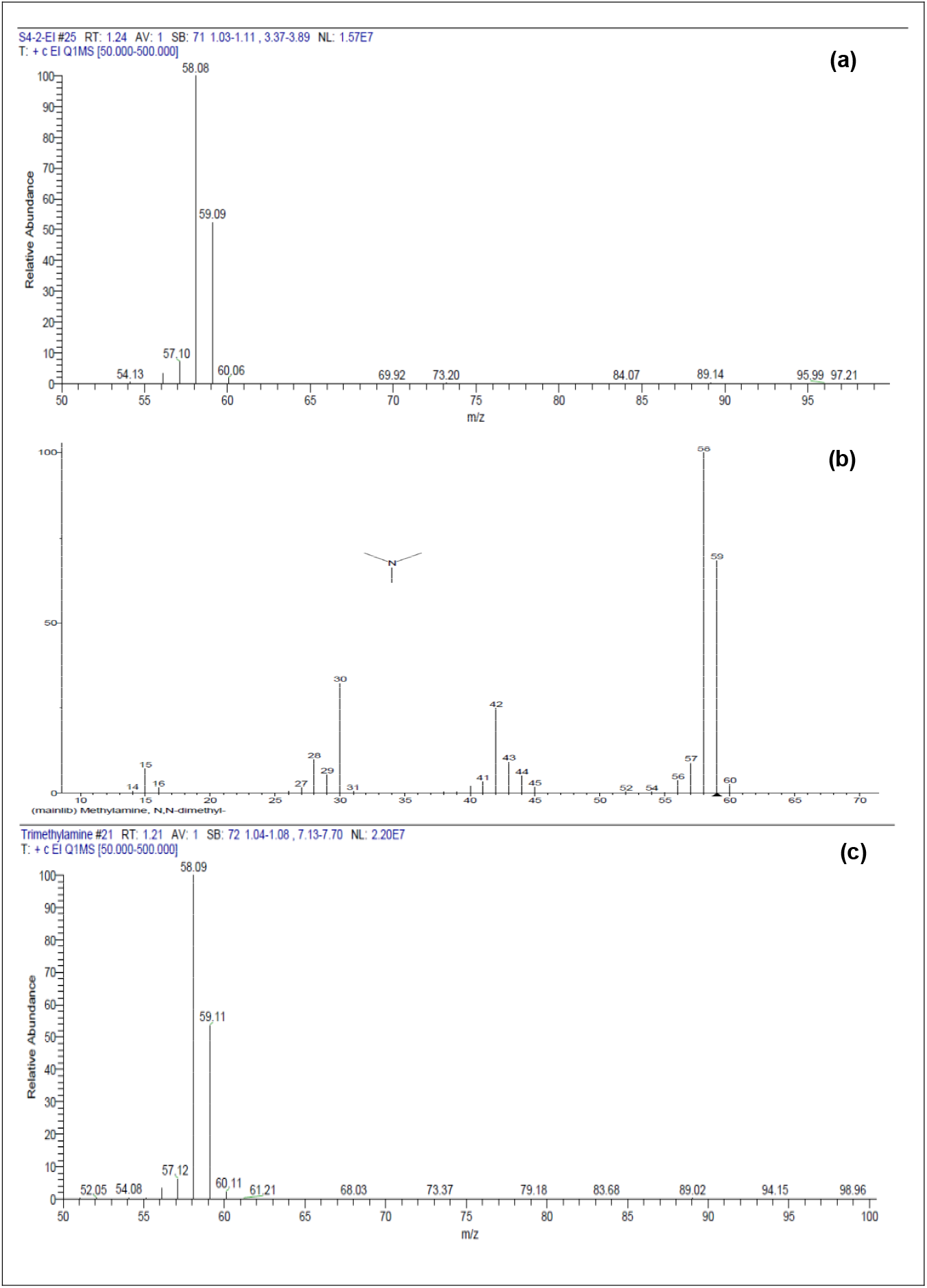
**(a)** EI-Mass spectra of trimethylamine at RT 1.4 min in the Toyobo buffer Cat # QRZ-101 **(b)** EI-Mass spectra of trimethylamine given by the NIST Library **(c)** Experimental EI-Mass spectra of trimethylamine standard.

**Figure 12:**
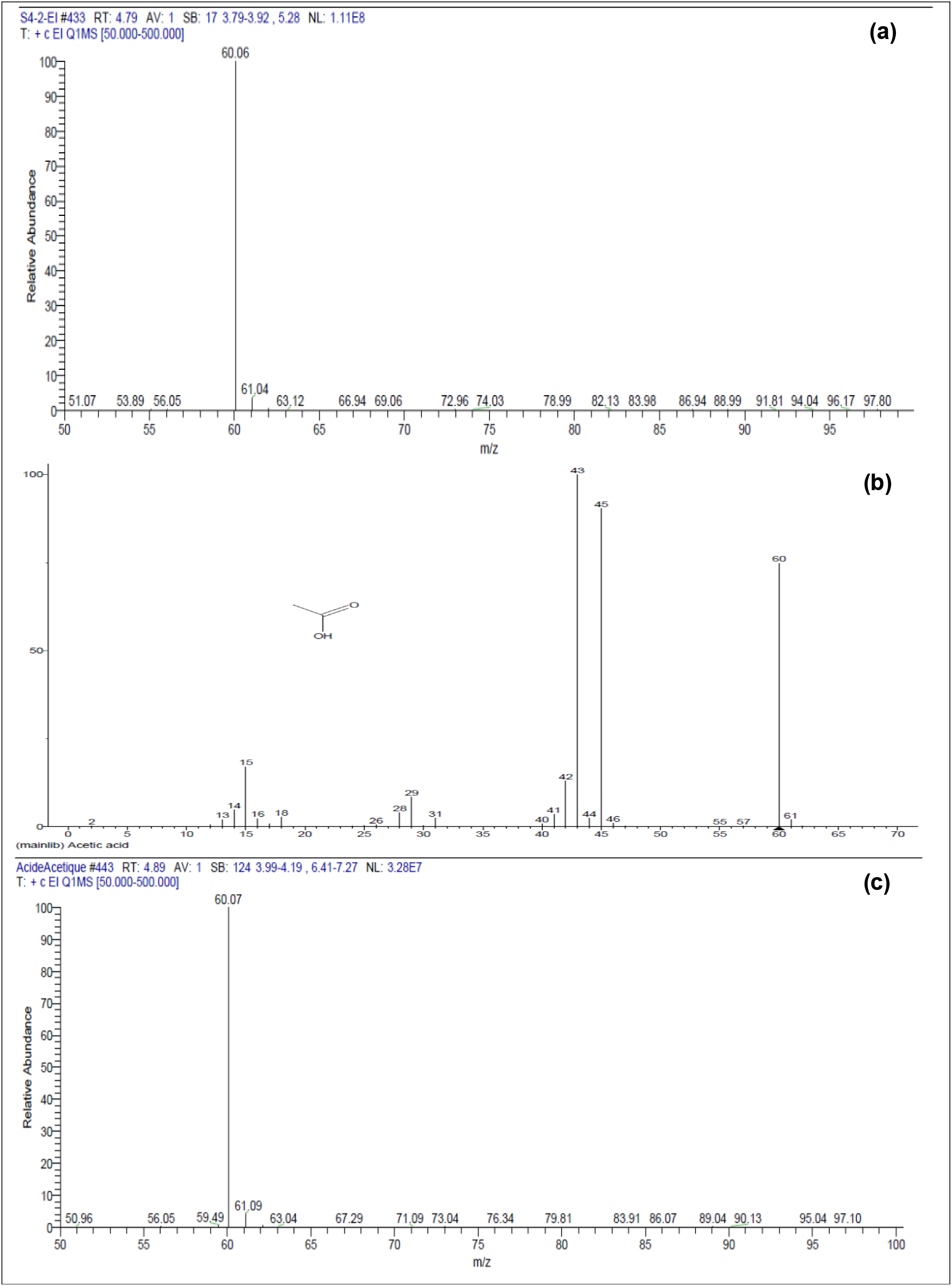
**(a)** EI-Mass spectra of acetic acid at RT 4.7 min in the Toyobo buffer Cat # QRZ-101 **(b)** EI-Mass spectra of acetic acid given by the NIST Library **(c)** Experimental EI-Mass spectra of acetic acid standard.

**Figure 13:**
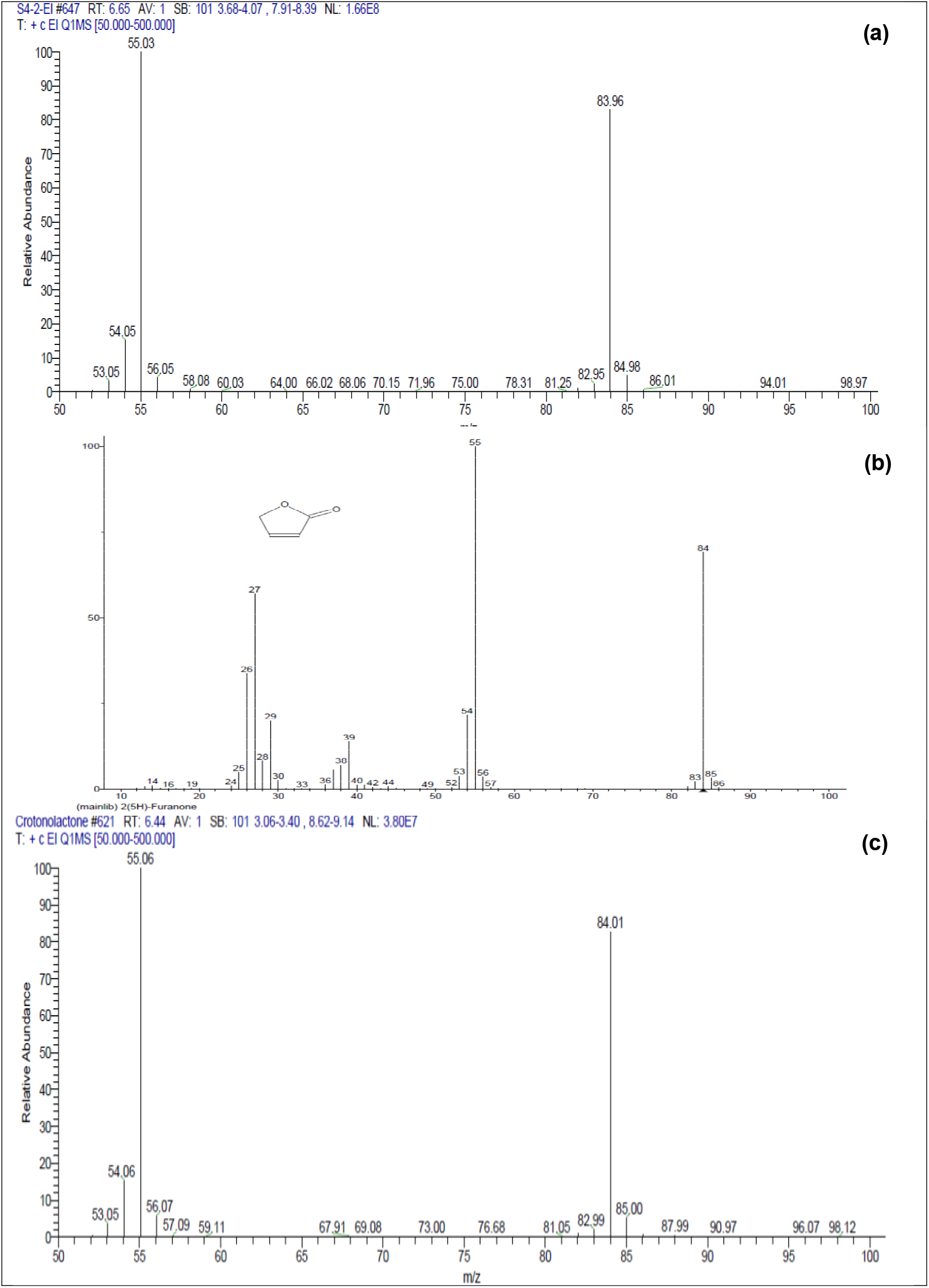
**(a)** EI-Mass spectra of crotonolactone at RT.6.5 min in the Toyobo Cat # QRZ-101 **(b)** EI-Mass spectra of crotonolactone given by the NIST Library **(c)** Experimental EI-Mass spectra of crotonolactone standard

**Figure 14:**
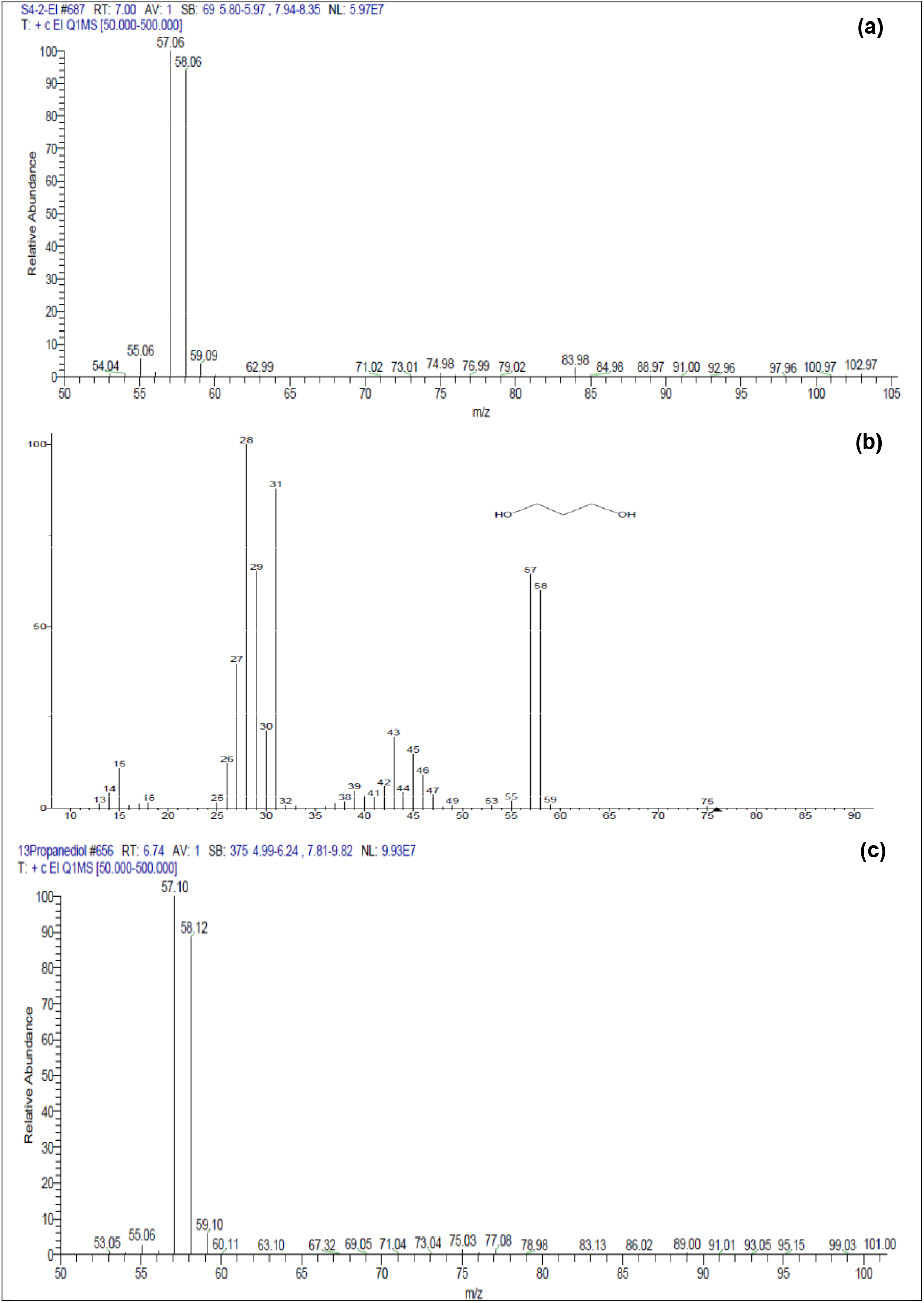
**(a)** EI-Mass spectra of 1,3-propanediol at RT 7.0 min in the Toyobo Cat # QRZ-101 **(b)** EI-Mass spectra of 1,3-propanediol given by the NIST Library **(c)** Experimental EI-Mass spectra of 1,3-propanediol standard

Overall, the chromatographic resolution of the method is very good as all the peaks are sufficiently separated from each other. Only peak # 3, identified as crotonolactone in the Toyobo buffer QRZ-101 (see Figure 1 in Appendix), appears less separated from the next peak # 4. The ability of the method to sensitively detect all co-solvents has been demonstrated as saturation peaks were observed. Eight co-solvents are presented in this list and each of them has been identified with a high level of confidence. The number of cosolvents is variable from one buffer provider to another. Some of them use only a small number of co-solvents such as DMSO alone or DMSO with some other such trimethylamine (TMA) or glycerol, or glycerol and isobutyramide. Others like Toyobo Company use more sophisticated buffers containing at leat 4 co-solvents such as TMA/acetic acid/1,3-propanediol/glycerol or crotonolactone.

DMSO is a frequently used co-solvent; it is identified in several PCR buffers. It is proposed to resolve secondary structure during PCR [15]. Isobutyramide binds in the major and minor groove of DNA and destabilizes the template double helix [2]. 1,3-Propanediol depresses the melting temperature of DNA [1]. Other co-solvents have been reported with similar potencies [3].

### 2.2 Co-solvent quantification

The co-solvents were quantified using MS operated in the single-ion-monitoring (SIM) which is the mode that offers the most sensitive and selective detection in most GC/MS method. Table 5 shows the levels of some co-solvents found in NEB and Toyobo buffers. The results were obtained in duplicate and show that the method has a good reproducibility. The accuracy of the method was determined by analyzing some buffers spiked with known amount of co-solvent. The spiking levels were close to 2.5 % and 4.5 %. Tables 6 and 7 illustrate the results obtained for a Toyobo buffer spiked with 1,3-propanediol and a NEB buffer spiked with DMSO, respectively. In both cases, the recovery was satisfactory in the range 90-120 %

**Table 5 :** Assay of Co-solvents in weight / weight percent

| Buffer<br>(Cat #) | Isobutyramide | 1,3-Propanediol | DMSO | Glycerol |
| --- | --- | --- | --- | --- |
| NEB<br>(M0491s), 5x | 4.5<br>4.6<br>**Mean: 4.6** |  | 15.3<br>14.9<br>**Mean: 15.1** | 25.1<br>25.9<br>**Mean: 25.5** |
| TOYOBO<br>(QPS-101), 2x |  | 11.7<br>12.0<br>**Mean: 11.9** |  |  |
| TOYOBO<br>(QPS-201), 2x |  | 11.9<br>12.3<br>**Mean: 12.1** |  | 2.6<br>2.6<br>**Mean: 2.6** |
| TOYOBO<br>(QRZ-101), 1x |  | 6.4<br>6.7<br>**Mean: 6.6** |  |  |
| TOYOBO<br>(QKD-201), 2x |  | 12.7<br>12.2<br>**Mean: 12.5** |  | 2.1<br>1.9<br>**Mean: 2.0** |

**Table 6:**
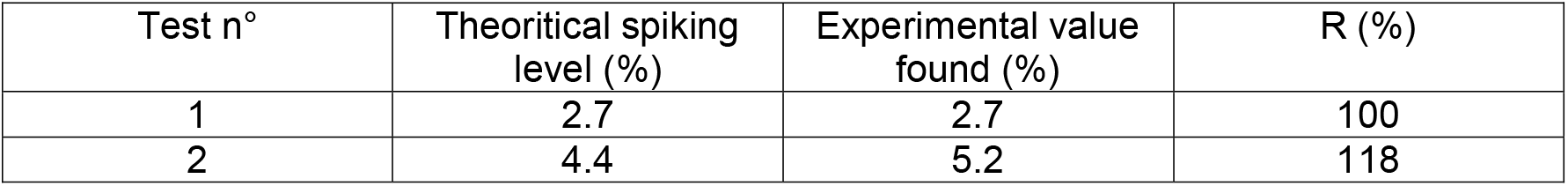
Recovery ratio of 1,3-propanediol in a Toyobo buffer

**Table 7:**
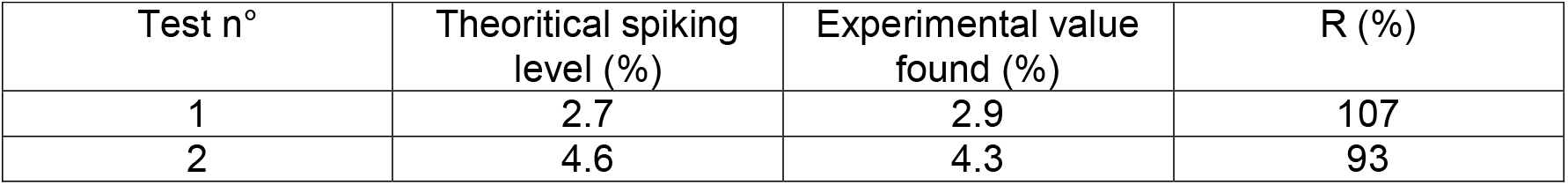
Recovery of DMSO in a NEB buffer

### 2.3. Biochemical evaluation of co-solvents

We assessed the amplification of a GC-rich template (c-Jun, 64% GC, 996 bp) by isobutyramide and 1,3-propanediol (Table 5). We also included 1,4-butanediol, a structural analog of 1,3-propanediol, for comparison. In the absence of co-solvent, no PCR product is detected. All three co-solvents amplify c-Jun to a similar extent (Figure 15).

**Figure 15.**
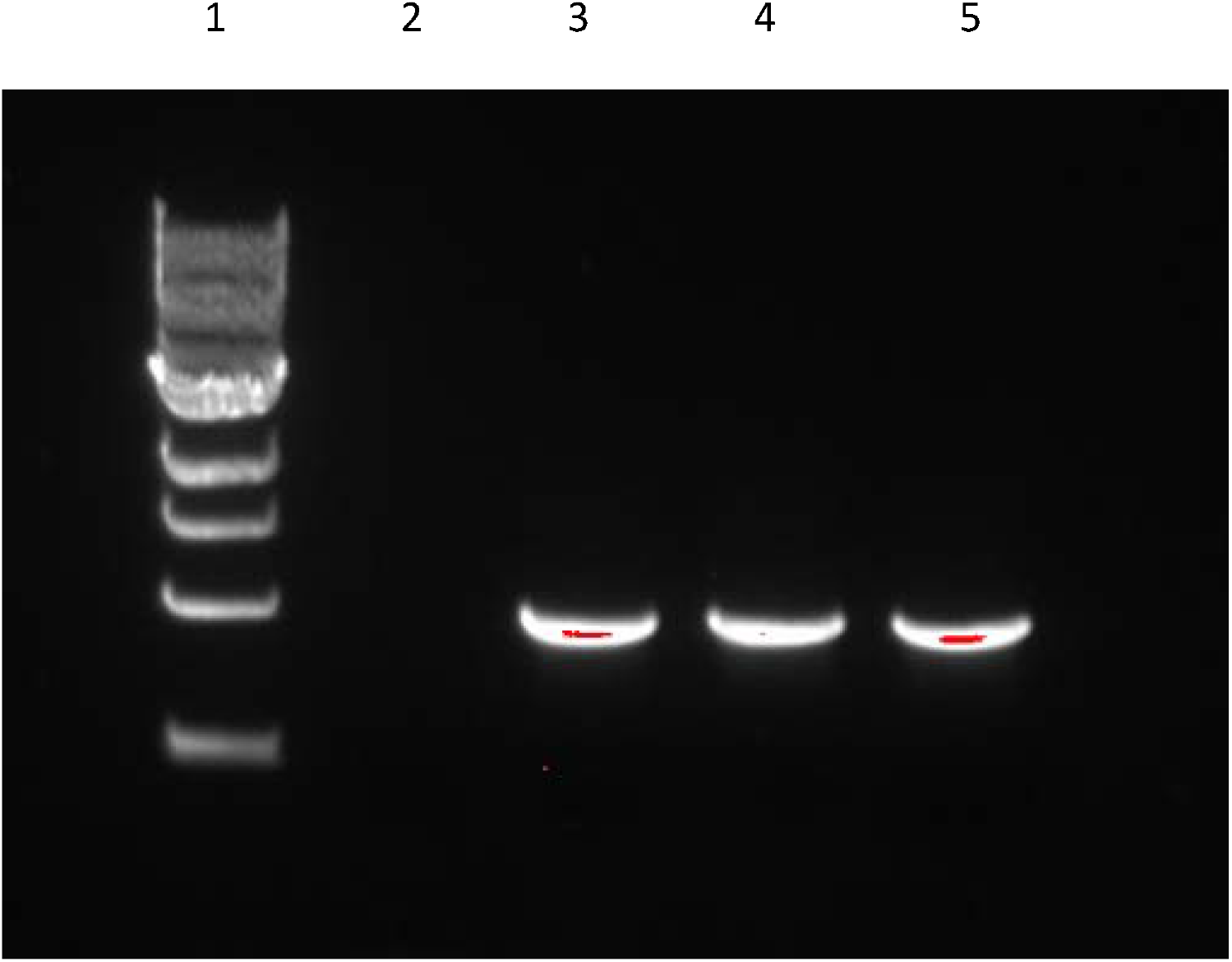
Effect of various cosolvents on c-jun template amplification. Lane 2: no additive; Lane 3: 0.9%isobutyramide; Lane 4: 6% 1,3-propanediol; Lane 5: 5% 1,4-butanediol.

We evaluated the effect of the same co-solvents on the thermostability and activity of lysates of wild type Taq. We used a range of concentrations of co-solvents (from one half to two-fold the concentrations listed in Table 5). The results are shown in Table 8. Thermostability in the presence of 0.9% isobutyramide and 6% 1,3-propanediol remains intact when compared with Taq that contains no additives. In the presence of 5% 1,4-butanediol, Taq retains only 35% of its thermostability; the Tm of the template decreases to 88.5°C from 92.5°C. At concentrations higher than those used in the commercial products, Taq thermostability is compromised. Inclusion of glycerol in the PCR buffer results in increases in Taq thermostability, presumably because glycerol stabilizes the protein. The effect of co-solvents on the screening activity score is generally similar to that of screening thermostability score, but note that the effect of glycerol in recovering activity is generally less than its effect in recovering thermostability, consistent with the primary effect of glycerol being on stability (Table 8).

**Table 8.** RT-PCR thermostability and activity of Taq lysates in the presence of co-solvents (N=2). Melting temperatures of DNA products at the end of PCR cycling are also presented.

|  | Taq thermostability |  | Taq activity |  |
| --- | --- | --- | --- | --- |
|  | peak area/cell # | Tm (°C) | peak area/cell # | Tm (°C) |
| no additives | 21.8 | 92.5 | 21.5 | 92.5 |
| 0.45% isobutyramide <sup>(1)</sup> | 22.0 | 91.5 | 21.7 | 91.5 |
| 0.9% isobutyramide | 19.9 | 91.0 | 20.2 | 91.0 |
| 0.9% isobutyramide, 5% glycerol | 21.0 | 91.0 | 21.8 | 91.0 |
| 1.35% isobutyramide | 16.0 | 91.0 | 19.9 | 91.0 |
| 1.35% isobutyramide, 7.5% glycerol | 21.7 | 91.0 | 21.7 | 90.0 |
| 1.8% isobutyramide | 10.9 | 90.0 | 11.5 | 90.5 |
| 1.8% isobutyramide, 10% glycerol | 17.4 | 90.0 | 15.1 | 90.0 |
| 3% 1,3-propanediol | 21.7 | 91.0 | 20.5 | 91.0 |
| 6% 1,3-propanediol | 20.8 | 90.0 | 21.1 | 91.0 |
| 6% 1,3-propanediol, 1.3% glycerol | 19.9 | 90.5 | 21.1 | 90.5 |
| 9% 1,3-propanediol | 17.3 | 90.0 | 17.1 | 90.0 |
| 9% 1,3-propanediol, 1.9% glycerol | 22.0 | 90.0 | 20.5 | 90.5 |
| 12% 1,3-propanediol | NA | NA | NA | NA |
| 12% 1,3-propanediol, 2.6% glycerol | 18.0 | 90.0 | 14.5 | 90.0 |
| 2.5% 1,4-butanediol | 21.6 | 91.0 | 21.4 | 91.0 |
| 5% 1,4-butanediol | 7.8 | 88.5 | 13.0 | 88.5 |
| 7.5% 1,4-butanediol | NA | NA | 7.9 | 88.0 |

<sup>(1)</sup> isobutyramide samples contained 2% DMSO.
NA: Not measured

We also screened cosolvents for thermostability and measured their effects on enzyme specific activity using purified Taq protein (Table 9). The results for thermostability are comparable to those of the lysates, while the activity assay measures enzyme specific activity independently of any PCR amplification.

**Table 9.**
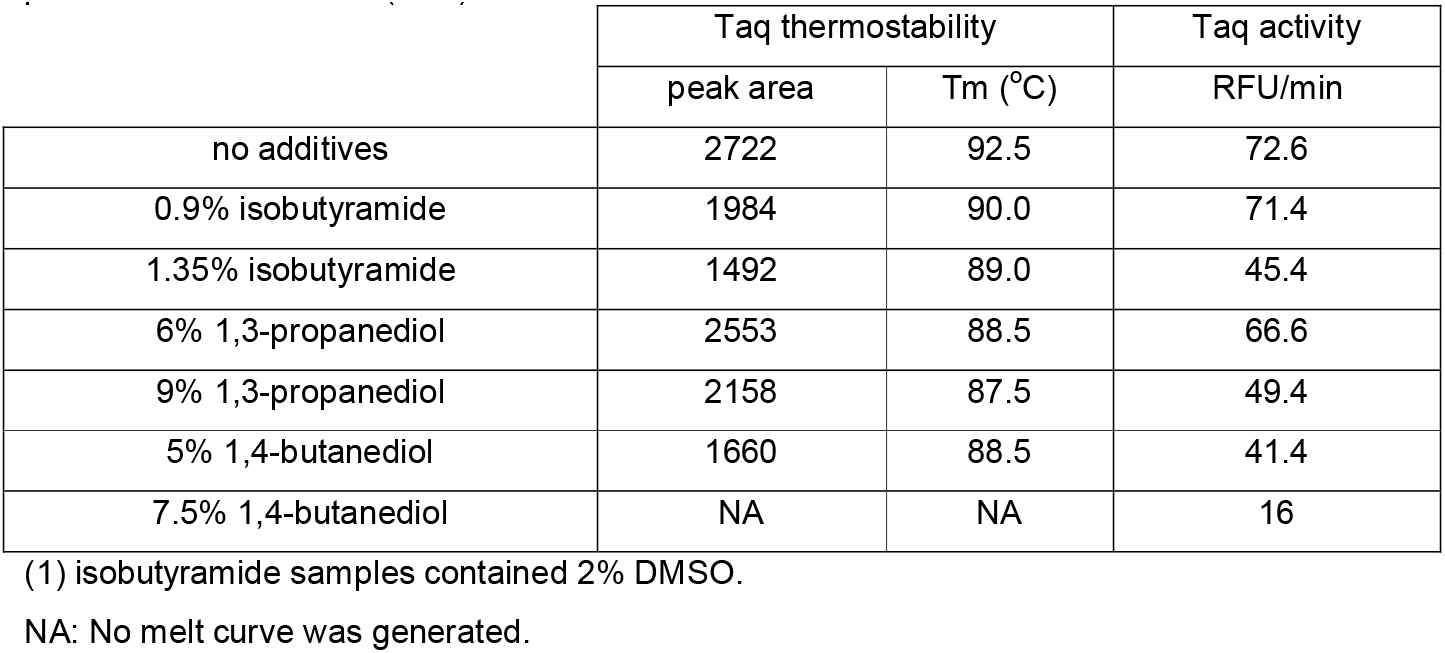
RT-PCR screening of thermostability and specific activity of purified Taq in the presence of co-solvents (N=2).

DNA melting curves for the WT lysates at the end of thermal cycling are shown in Fig. 16. In the absence of co-solvent, there is significant primer-dimer formation. Primer-dimers are eliminated in the presence of 6% 1,3-propanediol and 5% 1,4-butanediol. Addition of 0.9% isobutyramide leads to reduction of the primer-dimer peak.

**Figure 16.**
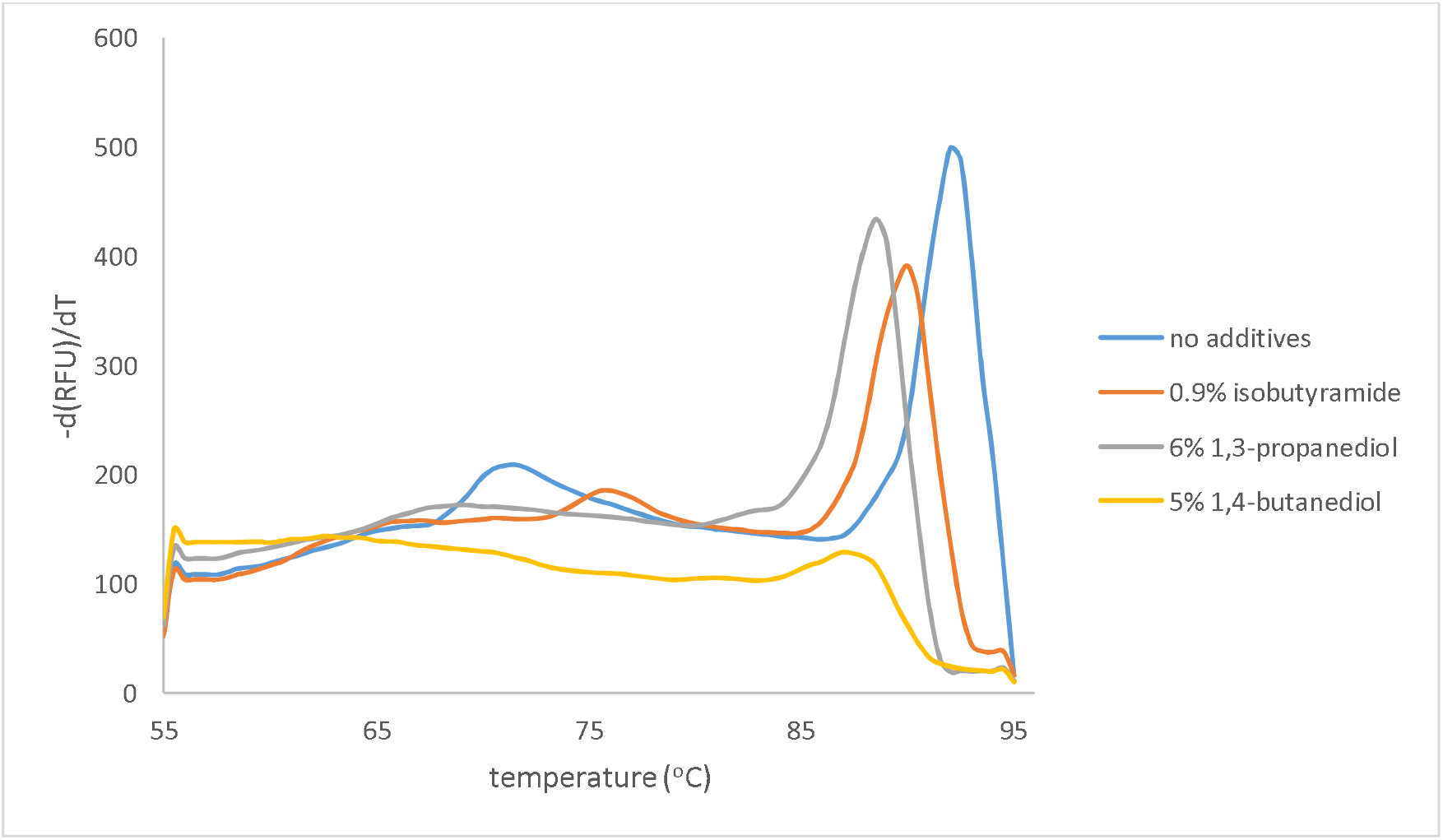
DNA melting curves for lysates of WT Taq in the absence and presence of co-solvents.

We also conducted thermal denaturation experiments in the presence of 5% 1,4-butanediol. The half-life of WT and T8 (a thermostable Taq carrying the mutations F73S, R205K, K219E, M236T, E434D, A608V) [18] at 95°C are 16 and 31 min, respectively. The presence of 5% 1,4-butanediol results in significantly reduced thermostability for both the WT and T8 (t_1/2_ 3.5 and 9.5 min, respectively).

It should be noted that 1,4-butanediol is a more effective co-solvent than 1,3-propanediol at least for certain templates with high GC content [1] It is probably due to the adverse effects of 1,4-butanediol on the polymerase (Tables 8,9) that Toyobo selected 1,3-propanediol as the co-solvent in their commercial products.

We note that the inclusion of isobutyramide as well as DMSO in the NEB Q5 mastermix shows that DMSO is not sufficient for robust amplification of GC-rich templates or for reducing GC-bias in next-generations sequencing applications. This is consistent with our prior work [1–3] which demonstrated the superior enhancement by higher molecular weight amides and sulfones compared to formamide and DMSO for a wide variety of GC-rich DNA templates.

## 3 Discussion

An analytical GC/MS method to identify and quantify the co-solvents in commercial aqueous PCR Buffers is presented. The buffers were first lyophilized and filtered for efficient removal of water and salified substance. The analytical capacity of the method was demonstrated by co-solvent analysis in 8 buffers from different manufacturers. All the major co-solvents were separated in less than 30 minutes, detected with a high sensitivity and identified with the mass spectroscopy detector. The use of deuterated co-solvent standards is a reliable technique for quantifying co-solvents in PCR buffers.

Properties of Taq polymerase in the presence of select co-solvents were evaluated. The identification and quantification of PCR enhancers are important to understand activities of commercial buffers and eventually improve their performance. The effects of cosolvents on the PCR amplification can be studied according to quantitative models for PCR amplification [1, 19]. These models show that the optimal co-solvent concentration corresponds to that where the favorable effects of the co-solvent on DNA melting and secondary structure outweigh the unfavorable effects on polymerase stability and activity with the greatest margin; also, the maximal cosolvent concentration compatible with PCR generally corresponds to that beyond which the polymerase activity is extinguished. While protein stabilizing agents like glycerol can partially compensate for the unfavorable effects of cosolvents on polymerases, as shown above they rescue polymerase activity to a lesser extent, and moreover, it is known that protein engineering is more effective in improving polymerase thermostability as well as reducing the detrimental effects of certain polymerase inhibitors [18]. As such, it is anticipated the polymerase engineering in the presence of polar organic co-solvents may enable the use of these compounds in significantly higher quantities and with greater enhancing effects. As such we are applying directed evolution to engineer a solvent-resistant Taq polymerase, in order to enable further exploitation of their favorable effects on PCR. For example, polymerase engineering may enable the use of 1,4-butanediol rather than 1,3 propanediol to achieve better DNA melting improvement while maintaining reasonable polymerase stability and acitivity, across a wide variety of applications. More generally, the types of compatible co-solvents and the maximum tolerated co-solvent concentrations can be improved by such methods. Finally, we note that the inclusion of cosolvents in so many leading mastermixes also shows they are recommended in nearly all applications and that such engineered Taq polymerases would therefore find very general utility in PCR.

## References

[1] R. Chakrabarti, Novel PCR-Enhancing Compounds and Their Modes of Action, in: T. Weissensteiner, HG Griffin, A. Griffin (Eds), PCR Technology Current Innovations. 2nd edition, CRC Press, 2003: pp. 51–63.

[2] R. Chakrabarti, C.E. Schutt, The enhancement of PCR amplification by low molecular weight amides, Nucleic Acids Res. 29 (2001) 2377–2381.

[3] R. Chakrabarti, C.E. Schutt., The enhancement of PCR amplification by low molecular weight sulfones, Gene 274 (2002) 293–298.

[4] D.D. Pratyush, S. Tiwari, A. Kumar, S.K. Singh, A new approach to touch down method using betaine as co-solvent for increased specificity and intensity of GC rich gene amplification, Gene 497 (2012) 269–272.

[5] K. Varadaraj, D.M. Skinner, Denaturants or co-solvents improve the specificity of PCR amplification of a G+C-rich DNA using genetically engineered DNA polymerases, Gene 140 (1994) 1–5.

[6] U.H. Fray, H.S. Bachman, J. Peters, W. Siffert, PCR-amplification of GC-rich regions: “slowdown PCR”, Nature Protocols 8 (2008) 1312–1317.

[7] A. Takeuchi, S. Yamamoto, R. Narai, M. Nishida, M. Yashiki, N. Sakui, A. Namera, Determination of dimethyl sulfoxide and dimethyl sulfone in urine by gas chromatography-mass spectrometry after preparation using 2,2-dimethoxypropane, Biomed Chromatogr. 24 (2010) 465–471.

[8] ASTM D6584-17, Standard Test Method for Determination of Total Monoglycerides, Total Diglycerides, Total Triglycerides, and Free and Total Glycerin in B-100 Biodiesel Methyl Esters by Gas Chromatography, ASTM International, West Conshohocken, PA, 2017.

[9] V. Gembus, J.P. Goull, C. Lacroix, Determination of glycols in biological specimens by gas chromatography-mass spectrometry, J. Anal. Toxicol., 26 (2002) 280–285.

[10] K.A. DaCosta, J.J. Vrbanac, S.H. Ziesel, The measurement of dimethylamine, trimethylamine, and trimethylamine N-oxide using capillary gas chromatography-mass spectrometry, Anal. Biochem. 187 (1990) 234–239.

[11] NIST mass spectral database http://webbook.nist.gov/ (accessed September 27, 2021)

[12] B. Yu, Y. Song, H. Yu, L. Huan, H. Liu, Optimizations of large volume-direct aqueous injection-gas chromatography to monitor volatile organic compounds in surface water, Analytical Methods 6 (2014) 6931–6938.

[13] C. Aeppli, M. Berg, T.B. Hofstetter, R. Kipfer, R.P. Schwarzenbach, Simultaneous quantification of polar and non-polar volatile organic compounds in water samples by direct aqueous injection-gas chromatography/mass spectrometry, Journal of Chromatography A 1181 (2008) 116–124.

[14] A.I. Dragan, R. Pavlovic, J.B. McGivney, J.R. Casas-Finet, E.S. Bishop, R.J. Strouse, M.A. Schenerman, C.D. Geddes, SYBR Green: fluorescence properties and interaction with DNA, J. Fluoresc. 22 (2012) 1189–1199.

[15] Y. Wang, D.E. Prosen, L. Mei, J.C. Sullivan, M. Finney, P.B. Vander Horn, A novel strategy to engineer DNA polymerases for enhanced processivity and improved performance in vitro, Nucleic Acids Res. 32 (2004) 1197–1207.

[16] F.C. Lawyer, S. Stoffel, R.K. Saiki, S.Y. Chang, P.A. Landre, R.D. Abramson, D.H. Gelfand, High-level expression, purification, and enzymatic characterization of full-length Thermus aquaticus DNA polymerase and a truncated form deficient in 5’ to 3’ exonuclease activity, PCR Methods Appl. 2 (1993) 275–287.

[17] J. Strien, J. Sanft, G. Mall, Enhancement of PCR amplification of moderate GC-containing and highly GC-rich DNA sequences, Mol. Biotechnol. 54 (2013) 1048–1054.

[18] F.J. Ghadessy, J.L. Ong, P. Holliger P, Directed evolution of polymerase function by compartmentalized self-replication, Proc. Natl. Acad. Sci. USA 98 (2001), 4552–4557.

[19] K. Marimuthu, C. Jing, R. Chakrabarti., Sequence-dependent biophysical modeling of DNA amplification, Biophys. J. 107 (2014) 1731–174.

